# Structural specialization of mossy fiber boutons is necessary for their unique computational functions

**DOI:** 10.1101/2025.10.20.683513

**Authors:** Nishant Singh, Suhita Nadkarni

## Abstract

Sparsely active granule cells in the dentate gyrus (DG) project onto CA3 pyramidal neurons through mossy fibers (MF), forming a feedforward network that ensures even similar inputs activate distinct and minimally overlapping populations of CA3 neurons. This process, known as pattern separation, is a key computational feature at this synapse. However, such sparse activity and connectivity increase the risk of information loss, as multiple presynaptic inputs are typically required to elicit a postsynaptic action potential. Interestingly, MF synapses exhibit robust short-term plasticity (STP), enabling a dynamic increase in release probability during brief bursts of presynaptic activity. This mechanism ensures that a short, high-frequency burst from a single granule cell can reliably generate a spike in its postsynaptic CA3 target. Unlike other hippocampal synapses, MF boutons are large, with multiple active zones, each coupled to a cluster of voltage-dependent calcium channels (VDCCs). MF boutons also possess a large readily releasable pool of vesicles. The functional consequences of this unusual synaptic design remain largely unexplored. In fact, the MF synapse is often depicted as a synapse with multiple sites, each behaving as an independent transmission line, analogous to several CA3 boutons contacting a single CA1 dendrite. We developed a physiologically realistic spatial model of the MF bouton to investigate how its peculiar structural and functional properties affect synaptic signaling and plasticity. Contrary to earlier assumptions, our computational model revealed, release sites are not independent transmission units. Crosstalk between calcium domains is necessary for the observed strong STP and for the timely activation of CA3 neurons. VDCCs in the MF bouton are only loosely coupled to active zones, and the distance between active zones is relatively large. In addition to the synaptic design and the known role of calbindin-D28k and synaptotagmin-7 in STP, we find that loose coupling of VDCCs to active zones and large inter-AZ distances are crucial. These features keep the basal release probability low, and their combined action is required to generate the facilitation that triggers postsynaptic action potentials in response to bursts filtering out non-informative dentate activity, and provides a strong rationale for the mossy fiber’s synaptic architecture.

## Introduction

The hippocampus plays an essential role in spatial navigation and the formation of episodic memories. Within the hippocampus, the dentate gyrus (DG) receives multimodal sensory inputs from the entorhinal cortex via the perforant pathway, which synapses onto granule cells. The spatio-temporal patterns of neuronal signals arriving at the dentate gyrus encode an animal’s experience (Leutgeb et al., 2007). Lesions in this area compromise working memory (Sasaki et al., 2018; Xavier and Costa, 2009) and spatial navigation (Ahmadi et al., 2025).

Granule cells send unmyelinated mossy fiber (MF) axons to the CA3 region. These axons terminate in three different types of boutons: large mossy fiber terminals (4–10 µm), filopodial extensions arising from mossy fiber boutons (0.5–2.0 µm), and small en passant varicosities (0.5–2.0 µm). The morphology of the large mossy fiber boutons — the subject of our investigation — stands out compared to other synapses of the hippocampus. These boutons form multiple contacts with the proximal apical dendrites of CA3 pyramidal (CA3p) neurons and exhibit a complex ultrastructural organization. The key structural features of MF boutons are well characterized. These include the number and spatial distribution of active zones, the size of the readily releasable pool (RRP), and the organization of associated calcium channels. MF boutons contain anywhere from a few to several dozen active zones, arranged across the synaptic surface with an average spacing of about 0.5 µm (Rollenhagen et al., 2007). Each active zone houses a variable number of RRP vesicles and is typically coupled to a cluster of voltage-dependent calcium channels (VDCCs), whose distances from the release sites can vary (Rollenhagen et al., 2007; Vyleta and Jonas, 2014). In sum, the DG gives rise to highly diverse and structurally complex MF boutons, each capable of exerting substantial influence on synaptic signaling. However, the precise ways in which these presynaptic design features shape synaptic function—and the quantitative extent of their impact—remain poorly understood.

Unlike the Schaffer collaterals (axon bundles from CA3 to CA1 region), where long-term potentiation (LTP) dominates functional modulation, the functional role of MF synapses is believed to be mediated primarily through short-term synaptic plasticity (Urban et al., 2001). Knockout mice with impaired LTP or LTD (without affecting STP) in mossy fibers showed unimpaired learning (Huang and Sternberg, 1995; Yokoi et al., 1996). In contrast, STP disruption in mossy fibers (via genetic and pharmacological manipulations) impaired spatial learning (Silva et al., 1992b,a; Mayford et al., 1995). Their capability to rapidly enhance neurotransmitter release rate, a form of STP, is fundamental to critical functions at this synapse (Salin et al., 1996; Nicoll and Schmitz, 2005; Urban et al., 2001).

Despite the large number of release sites and RRPs, the probability that any vesicle is released in response to an action potential—release probability (Pr)—is surprisingly low at about 20% (Von Kitzing et al., 1994; Salin et al., 1996; Lawrence et al., 2004; Vyleta and Jonas, 2014; Chamberland et al., 2014). Consequently, a presynaptic AP in the mossy fiber is insufficient to elicit postsynaptic depolarisation strong enough to trigger CA3 pyramidal neurons. However, the MF-CA3p synapses show striking presynaptic short-term plasticity, consistent with their functional relevance. This swift escalation in neurotransmitter release rate in response to a brief, high-frequency stimulus can trigger an action potential in the postsynaptic CA3p neuron and is critical for encoding, storage, and recall of memory in the CA3 network (Vyleta and Jonas, 2014).

Although previous studies have shown that synaptotagmin-7 (the calcium sensor for asynchronous vesicle release) and saturation of the intrinsic calcium buffer contribute to STP (Jackman et al., 2016; Blatow et al., 2003; Matveev et al., 2004), the role of the remarkable design complexity of these terminals has not been explored in detail. In fact, most studies treat MFs as equivalent to multiple CA3 terminals; each active zone in the MF is regarded as an autonomous signal transmission line (Chamberland et al., 2018).

We examine the relationship between the intricate ultrastructure of the mossy fiber boutons (MFB) and the pronounced short-term plasticity characteristic of this synapse. If one considers the observed ranges of synaptic parameters, the resulting degeneracy in synaptic design would be enormous. This raises two key questions: are all combinations of these parameters physiologically viable, and what is the functional value of having a wide variety of possible synaptic configurations?

Our modeling framework offers an unprecedented platform for quantifying how the spatial organization of molecular components within the mossy fiber synapses influences information processing in the hippocampus. It also allows us to systematically investigate fundamental questions about form–function relationships and the impact of each physiologically plausible synaptic design on transmission and plasticity.

## Results

### Model Description

The computational model of the mossy fiber bouton consists of active zones, VDCCs, calcium ions, calbindin-D28k, PMCA pumps, and calcium sensors: synaptotagmin-1 (syt1) and synaptotagmin-7 (syt7). These molecular components, critical for triggering vesicle release, are well-organized within the synapse (Rollenhagen et al., 2007). We hypothesize that this elaborate arrangement is not merely incidental but influences synaptic transmissions and plasticity. Figure 1a illustrates an idealized mossy fiber synapse used in our simulations.

**Figure 1:**
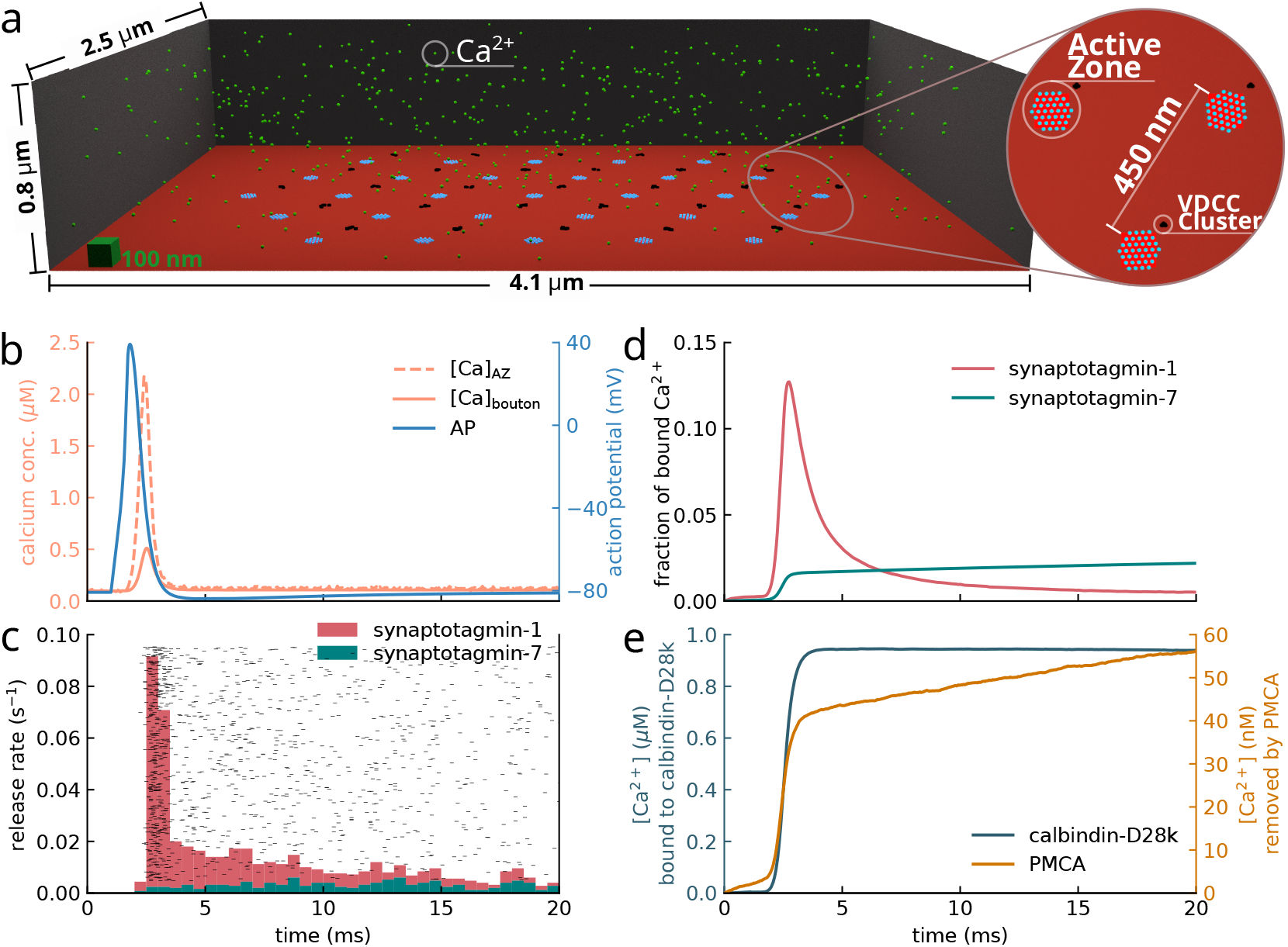
Mossy fiber bouton model. **(a)** An idealized 3D model of the mossy fiber bouton. Scale: 100 nm cube at bottom left. The zoomed-in portion of the white circle illustrates the spatial organization of the AZs and VDCC clusters. **(b)** An illustrative action potential and subsequent change in calcium concentration at the active zone and in the whole bouton. **(c)** Corresponding vesicle releases from 500 trials and vesicle release rate. **(d)** Fraction of bound calcium in syt1 and syt7. **(e)** Capture and extrusion of calcium ions by calbindin-D28k and PMCA pumps, respectively.

The MFB model has a volume of 8.2 µm^3^. Its plasma membrane, which borders the synaptic cleft, has multiple active zones, mutually separated by 450 nm (Rollenhagen et al., 2007). Each active zone has a coupled cluster of VDCCs and hosts several vesicles in its readily-releasable pool (Figure 1a). As action potential arrives at this bouton, calcium ions rush into the cytosol through VDCCs and transiently increase local and global calcium concentration (Figure 1b). These ions can bind to calcium sensors (Figure 1c). When all the calcium-binding sites of the sensors have captured a calcium ion (synaptotagmin-7 has five, and synaptotagmin-1 has two calcium-binding sites), neurotransmitter molecules are released into the synaptic cleft. Synaptotagmin-1 has a low affinity for calcium ions but a fast binding rate, and therefore, has the highest probability of binding during the peak of calcium influx, and therefore, mediates synchronous neurotransmitter release. Synaptotagmin-7, on the other hand, has distinct biophysical properties compared to syt1. Despite slower binding rates, the high calcium-binding affinity of this sensor ensures release at lower calcium concentrations. Consequently, the synchronous release rate is high but shortlived, and the asynchronous release rate is low but lasts longer (Figure 1d). Calcium buffer, calbindin-D28k, quickly binds a large fraction of the incoming calcium flux through VDCCs, while PMCAs slowly pump out extra calcium ions into the extracellular space, thereby regulating the free calcium inside the bouton (Figure 1e).

### Vesicle release probability of MFB designs

In our model, we simulate up to 15 calcium channels in a single cluster, 7-35 active zones, and 5-60 RRP (Rollenhagen et al., 2007). The maximum coupling distance possible between an AZ and its VDCC cluster (halfway between two adjacent active zones) was 220 nm. Consequently, we created 24,300 MFB designs with these four varying attributes summarised in Table 1. Each combination of these attributes gives a different bouton design. Are all these possible synaptic designs physiologically viable? What functional advantage might this large variability in design provide? To address these questions, each configuration was systematically stimulated with an action potential across multiple trials.

**Table 1.**
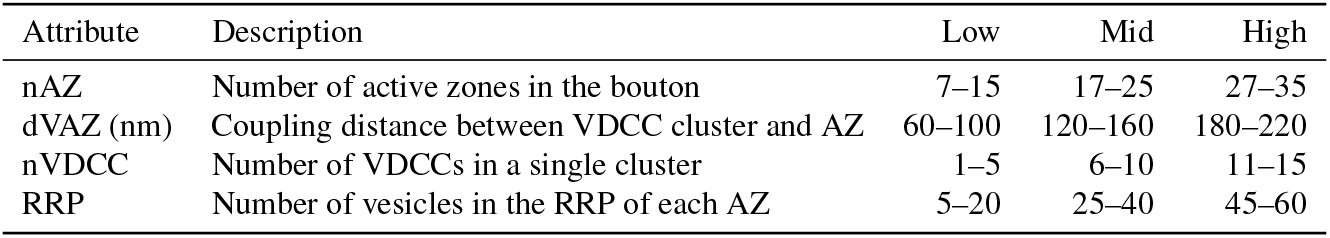
Synaptic attributes.

Within the parameter space, our simulations show that only a subset of designs support the physiological range of Pr: 5 % of all the designs have physiological release probability in the range 0.20 to 0.28 (Figure 2a). Among the designs that produce physiological levels of Pr, a notable characteristic is their bias toward larger values of nVDCC, dVAZ, and RRP. One interpretation of these results is that a large number of channels in close proximity to the active zone would lead to unphysiologically high release probabilities. Consequently, when channel numbers are high, their spatial positions must be constrained farther from the active zone to maintain a low release probability.

**Figure 2:**
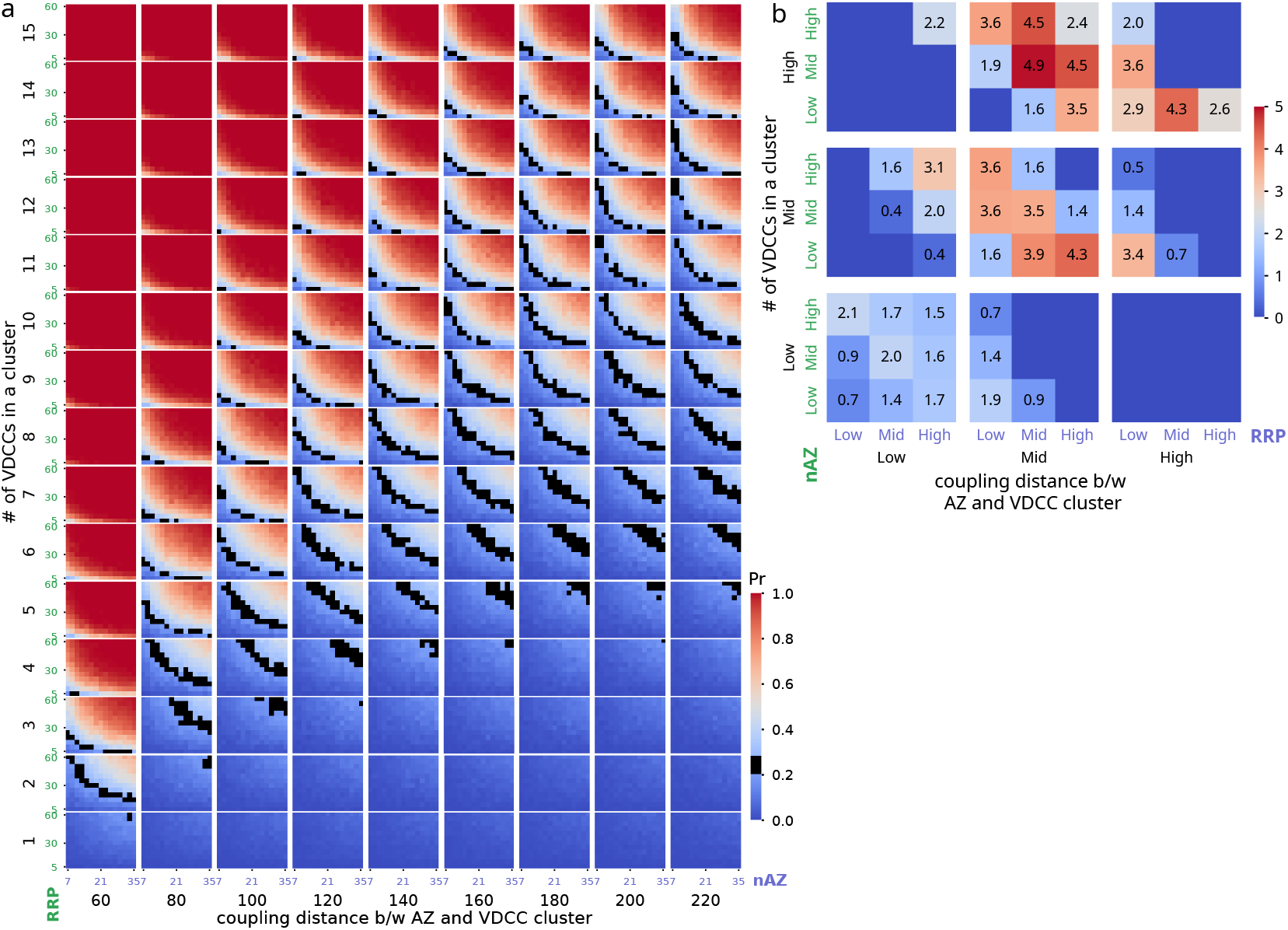
Release probability of MFBs. **(a)** Release probability of MFB designs with different synaptic attributes: number of VDCCs in a cluster, coupling distance between AZ and their VDCC cluster, number of AZ, and RRP size. Black spots indicate designs with Pr within 0.20 - 0.28. **(b)** Each parameter range was divided into three categories: low, mid, and high. The percentage of designs with physiological Pr (within 0.20 - 0.28) that fall in a category is shown.

To further dissect the complexity of the wide range of synaptic attributes—and to understand how these parameters collapse as additional experimental constraints are applied, as well as to explore the relationships among these highly non-linear variables—we divided each attribute into three levels (‘low’, ‘mid’, and ‘high’), resulting in 81 distinct categories of MFB designs (Table 1). Figure 2b shows the fraction of designs that belong to each category. A large number of VDCCs that are closely associated with AZs have higher Pr, while the other extreme has very low Pr. Only 44 of the 81 categories contain designs with physiological Pr.

### Properties of the control boutons

We then investigated how synaptic dynamics are shaped by diverse synaptic designs. From 44 categories, we selected one representative physiological design each, all exhibiting low basal Pr. A train of ten action potentials at 50 Hz, typical of burst activity, rapidly elevates Pr, with nearly all designs saturating above 0.9 due to multiple active zones (Figure 3a). Consequently, the Pr_*n*_/Pr_1_ ratio poorly reflects short-term facilitation at MF-CA3p synapses, as it overlooks multivesicular release impacting postsynaptic responses.

**Figure 3:**
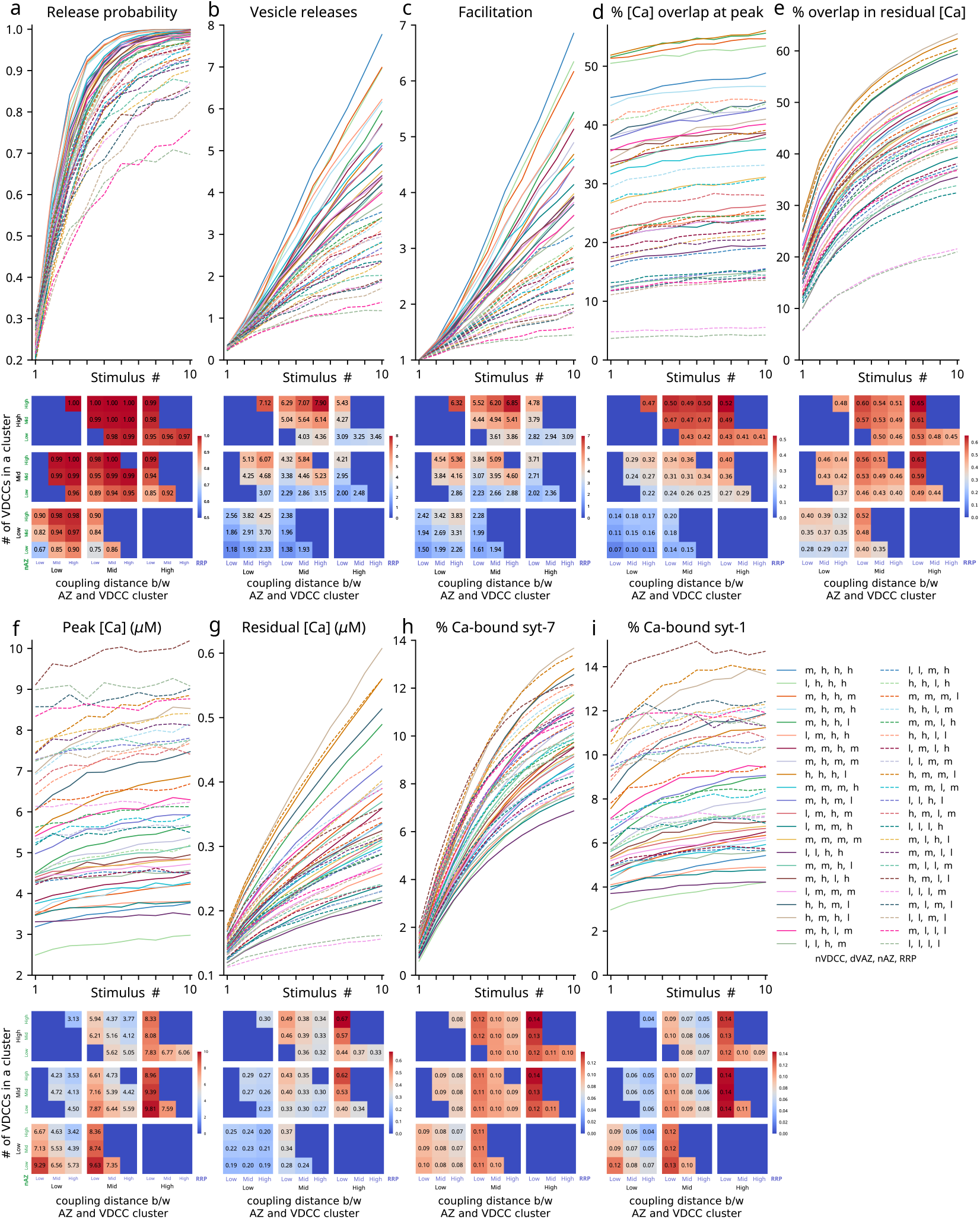
Properties of control boutons. We describe here the various synaptic properties of 44 design categories in response to a stimulus of ten APs at 50 Hz. The top row in each figure shows the response to the individual input spikes, while the heatmaps (below the line plots) show the distribution of these properties across designs in response to the tenth AP. **(a)** Release probability. **(b)** Number of vesicles released in response to an AP. **(c)** Facilitation. **(d)** Percentage of average calcium overlap at an active zone during an AP. **(e)** Percentage of average residual calcium overlap at the active zone. **(f)** Peak calcium concentration at the active zone after each AP. **(g)** Residual calcium concentration measured as the average concentration during 5-15 ms after an AP. **(h)** Fraction of occupied calcium binding sites in syt7. **(i)** Fraction of occupied calcium binding sites in syt1. Legend: l, m, and h represent ‘low’, ‘mid’, and ‘high’. Each combination containing four of these symbols (in order: nVDCC, dVAZ, nAZ, RRP) represents a design category.

The average number of vesicles released (⟨Rel_*i*_⟩) continues to increase with successive stimuli (Figure 3b). Therefore, to quantify facilitation, we use ⟨Rel_*i*_⟩ normalized by Pr_*i*_, which provides a more informative measure (Equation 1). This normalization mitigates large or spurious variations that could result from small initial values of ⟨Rel_*i*_⟩. Given the strong dependence of facilitation on Pr, this approach allows us to examine how facilitation (F_*i*_) varies for a fixed Pr across a train of stimuli.

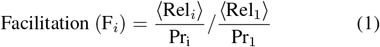

A subset of synaptic designs with fewer total VDCCs (nVDCC × nAZ) exhibits lower Pr saturation, with 13% showing Pr_10_ *<* 0.9 and 5% below 0.8, typically due to a smaller RRP (Figure 3a). Conversely, designs with numerous active zones and larger AZ-VDCC coupling distances exhibit the greatest increase in vesicle release (Figure 3b). Synaptic design significantly influences facilitation: shorter AZ-VDCC coupling distances suppress short-term plasticity, while larger RRPs or AZs enhance it. The highest STP levels occur in designs with large RRPs, extended coupling distances with many active zones, and moderate VDCC counts (Figure 3c).

We further probed facilitation, exploring additional mechanisms governing this process. During the calcium peak, the contribution of calcium from neighboring active zones to a site’s total calcium concentration grows with repeated stimulation (Figure 3d). Although residual calcium overlap is initially low, it increases sharply with successive stimuli (Figure 3e). Synaptic designs show varied degrees of overlap. Larger coupling distance favors a larger overlap. Increase in VDCCs and active zones also promotes this overlap (Figure 3d, e).

To quantify calcium spread from each VDCC cluster and its invasion of neighboring active zones, we assigned a unique calcium species to each cluster. This enabled precise tracking of diffusion within the terminal. With successive stimuli, contributions from nearby active zones to both peak and residual calcium concentrations increased. Peak calcium rises modestly, while residual calcium increases substantially, with the magnitude varying across design categories. Larger coupling distances and more active zones enhance residual calcium increases (Figure 3f, g).

Calcium sensors on synaptic vesicles—Synaptotagmins 7 and 1—bind incoming calcium ions with distinct kinetics. Syt7 progressively accumulates and retains calcium, while Syt1 shows modest increases per action potential. Synaptic design strongly influences the fraction of calcium from neighboring VDCC clusters reaching an active zone, resulting in varied calcium binding to synaptotagmins. Larger coupling distances enhance the percentage of calcium bound to synaptotagmins (Figure 3h, i).

### Mechanism of STP at mossy fiber boutons

When an action potential briefly opens VDCCs at the presynaptic bouton, a large amount of calcium ions transiently flows into the cytoplasm. Due to the high cytosolic concentration and fast binding kinetics of calbindin-D28k, most of the incoming calcium is quickly bound, effectively limiting both the magnitude and spatial spread of the calcium signal. In this scenario, it may be appropriate to describe the action of each active zone as autonomous, governed solely by the cluster of VDCCs associated with it. However, when the bouton receives a train of stimuli, this simplification no longer holds. Repeated calcium influx with successive stimuli locally reduces calbindin’s buffering capacity, as its calcium-binding sites become increasingly saturated. As a result, each calcium signal is stronger and travels farther than the previous one (Figure 4a, b). In this synapse, which contains multiple active zones, each coupled with a cluster of VDCCs, we consider the calcium domain associated with each individual set of VDCCs and active zones. As the domain size increases with stimuli, their overlap also increases (Figure 4c, d). Residual calcium conccentration shows a drastic increase in contribution from neighboring AZs, while the peak calcium concentration only moderately increases (Figure 4d). This “collaboration” further leads to an activity-dependent increase in the calcium signal at the active zones (Figure 4e). These findings point to a significant interplay between calcium buffering and inter–AZ crosstalk in shaping short-term plasticity, beyond the previously proposed role of synaptotagmin-7 alone (Jackman et al., 2016). To disentangle the contributions of these components and precisely quantify the underlying pronounced short-term plasticity observed in our system, we designed three distinct simulation conditions besides Control.

**Figure 4:**
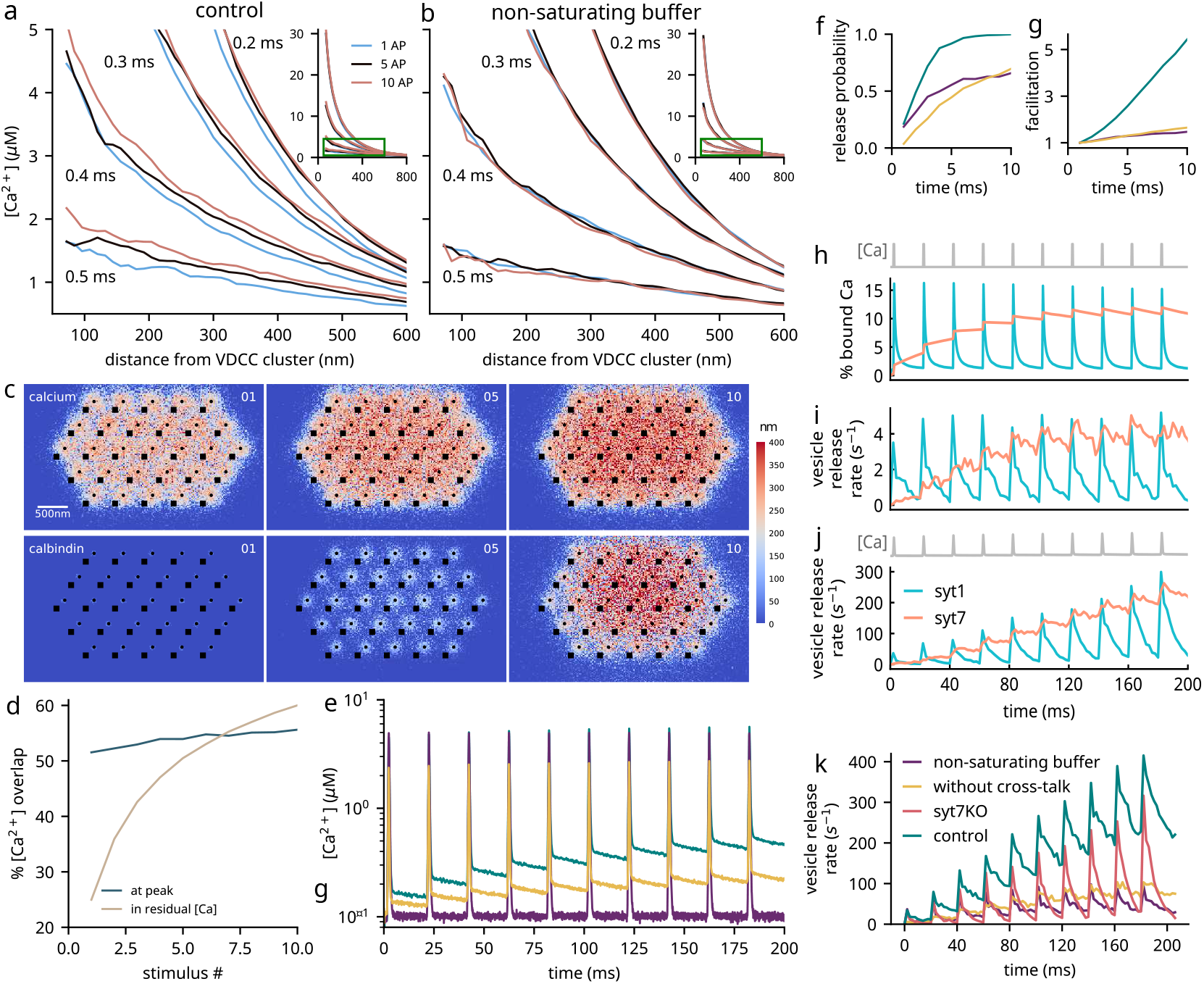
Mechanism of facilitation. **(a)** Calcium nanodomain enlarges with repeated stimuli. Calcium concentration decays with distance from a single cluster of VDCCs (measured at four timepoints) after the calcium concentration peaks in a control bouton. For each timepoint, three curves are shown corresponding to the 1st, 5th, and 10th AP. With each AP, these curves shift to the right, indicating an increase in the average distance calcium ions travel. **(b)** The buffering capacity of calbindins has been kept constant. As a result, there is no change in the spatial decay of calcium concentration. **(c)** In synapses with multiple active zones, the calcium nanodomains grow with multiple stimuli and contribute to other nearby active zones. (top row) The heatmap shows the height of the surface below which the calcium concentration exceeds 2 µM during an AP. (bottom row) The heatmap shows the surface height (in nm) below which at least 20% of calcium-binding sites in the buffer are occupied during an AP. The three heatmaps correspond to 1st, 5th, and 10th AP, respectively, as indicated by numerals at the top right. **(d)** Average % contribution of calcium ions at an active zone from VDCC clusters of other AZs increases with stimuli. The two curves show the overlap at the peak calcium concentration and the residual calcium response to 10 APs at 50 Hz. **(e)** Average calcium concentration at the active zones rises with each subsequent AP. **(f)** Release probability and **(g)** facilitation for different simulation conditions. With unsaturating buffers, calcium domains remain the same and do not increase their contribution to nearby active zones. Consequently, release probability and facilitation are reduced compared to the control. The same is observed when cross-talk between the active zones is restricted. **(h)** Average calcium ions bound to syt1 and syt7, and **(i)** vesicle release rate via syt1 and syt7 in response to an artificial calcium stimulus with peak height of 8 µM and a residual 300 nM. **(j)** Vesicle release rate via syt1 and syt7 in response to calcium influx from VDCCs in a realistic design. In addition to buffering and cross-talk, syt7 retains bound calcium ions in response to subsequent stimuli, leading to a sustained increase in vesicle release. Syt1, on the other hand, loses its calcium ions and contributes minimally to the increase in vesicle release. **(k)** Vesicle release rates in the four simulation conditions.

1. **No Crosstalk**: Each active zone responds only to calcium influx from its own local VDCC cluster, eliminating the role of inter-AZ calcium diffusion (to isolate the effect of overlap of calcium domains).
2. **Non-saturating Buffer (nonsatBuffer)**: Calcium buffering is modeled with a constant, non-depleting buffer concentration (to isolate effects of buffer saturation).
3. **syt7 Knockout (syt7KO)**: Synaptotagmin-7–mediated neurotransmitter release is abolished.

To simulate the synapse with fully autonomous calcium microdomains, we assigned unique calcium species to each active zone, allowing it to bind selectively to its “own” synaptotagmins. This prevented calcium ions released through a VDCC cluster from influencing sensors at neighboring active zones. As a result, local calcium concentrations at each AZ dropped sharply (Figure 4e). This intervention, feasible due to our morphologically detailed computational model, disrupts the spatial integration of calcium signals across neighboring active zones. This autonomy of calcium microdomains led to a striking reduction in vesicle release probability and facilitation (Figure 4f, g). These results highlight the critical role of calcium domain crosstalk in synaptic transmission. Far from being incidental, the spatial overlap of calcium signals across active zones emerges as a key determinant of short-term plasticity, underscoring the importance of coordinated calcium dynamics across microdomains and, morphological complexity of the synapse in synaptic function.

We simulate biophysically non-saturating buffers by keeping their buffering capacity unaffected by calcium influx. This is achieved by keeping the amount of available calcium-binding sites unaffected by calcium in the model. In this case, the calcium transient reaches a peak and drops back to its original baseline (100 nM). No residual calcium is observed. This trend remains constant over all subsequent stimuli (Figure 4e). Again, both synaptic transmission and short-term plasticity are severely compromised (Figure 4f, g).

Role of syt7 in STP at the MF-CA3p synapse has been reported (Jackman et al., 2016). Here, we explored the roles of both syt1 and syt7 as sensors for synchronous and asynchronous releases, respectively. We stimulated the model with a calcium signal containing a series of ten artificial square pulses. This eliminated the effect of mechanisms that enhanced calcium signals with subsequent stimuli to disambiguate the effect of only synaptotagmins on vesicle release. Vesicle release through syt1 in isolation closely follows high calcium transients. Vesicle release from syt7 shows slow calcium unbinding and, therefore, the subsequent increase in vesicle release rate even as the calcium signal remains the same after each stimulus (Figure 4h, i). In the natural case, the effect of buffers and overlapping calcium domains increases peak and residual calcium concentration. When the model is stimulated with a train of APs. This results in a subsequent increase in vesicle release via a synchronous pathway, which further augments facilitation by syt7. As a result, even in the syt7KO condition, small residual facilitation is observed (Figure 4j).

Each of the previously described mechanisms indeed does contribute to short-term plasticity, but remarkably, it is only when all molecular components act in concert under physiological conditions that we observe the full expression of facilitation consistent with experimental observation. This highlights a critical systems-level insight: high facilitation is not driven by any single factor alone, but emerges from the synergistic interplay of calcium buffering, VDCC distribution, synaptotagmin signaling, and inter-AZ crosstalk; each active zone does not act autonmously. Upon high-frequency stimulation, a non-linear amplification emerges: initially, a large fraction of calcium influx from VDCCs is rapidly sequestered by local buffers, restricting the spatial spread of calcium. However, with each successive spike, available buffer sites near VDCCs subsequently decreases, allowing calcium to diffuse more broadly. This progressive expansion of calcium spread leads to greater overlap between calcium domains from adjacent VDCC clusters (Figure 4d), dramatically elevating free calcium concentrations around active zones. Consequently, while calcium peak amplitudes increase modestly, residual calcium levels rise steeply across pulses (Figure 4e), driving robust facilitation as vesicle release increases (Figure 4h). These results reveal a powerful self-reinforcing mechanism: local reduction in buffering capacity and spatial calcium integration together shape a synapse highly tuned for short-term enhancement of release.

We next probed the effect of different designs on facilitation. Each of the 44 representative designs was simulated in all the simulation conditions. In most control designs, Pr increases quickly and saturates near the value 1. The next three cases—without crosstalk, nonsatBuffer, and syt7KO—show different degrees of reduction in Pr compared to control (Figure 5a). As expected, each defect reduces the release probability after ten spikes (Figure 5d). The true extent of loss of function is seen in the number of vesicles released in response to an action potential (Figure 5b, e). We also see a similar trend in facilitation (Figure 5c, f).

**Figure 5:**
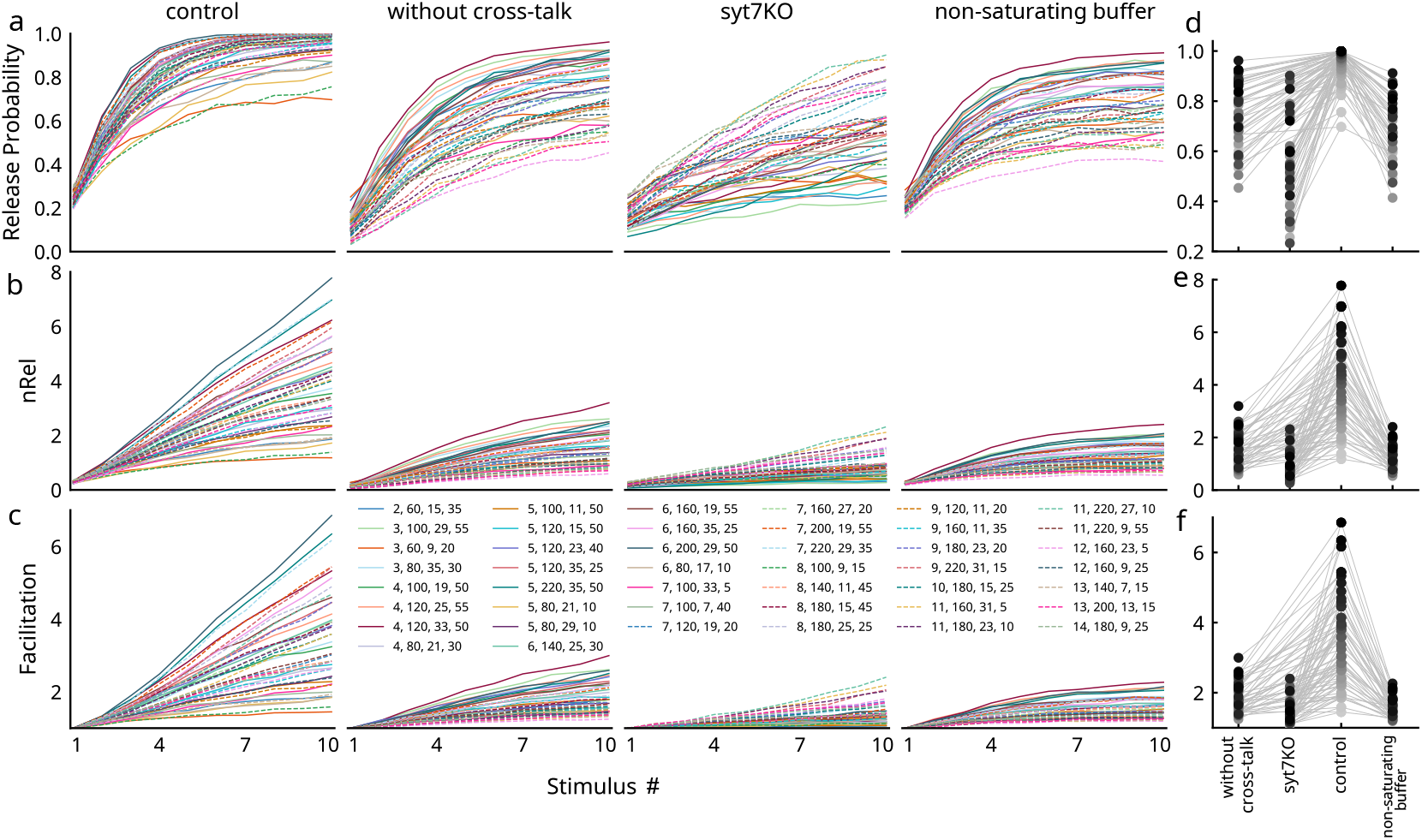
Facilitation in mossy fiber synaptic configurations. **(a)** Release probability in response to 10 APs at 50 Hz. Most control designs saturate close to a Pr of 1. In all the other three cases (without cross-talk, syt7KO, and unsaturating buffer) Pr is diminished by different amount for all the designs. **(b)** summarizes this reduction compared to control at the end of the 10th spike. **(c)** Compared to control, other cases see a drastic reduction in the total number of vesicles released after each AP. The response after 10th AP is summariszed in **(d). (e)** Similar trend is seen in facilitation: drastically low facilitation when cross-talk is restricted, syt7 is removed, or when buffers don’t saturate, summarized in **(f)**.

In general, higher values of nVDCC, dVAZ, and RRP in the control, no-crosstalk, and syt7KO conditions are consistently associated with enhanced facilitation. In contrast, the nonsatBuffer condition favored shorter coupling distances between active zones and VDCC clusters for enhanced facilitation, as greater separation limits calcium signaling. Notably, the design categories that contain a high proportion of physiologically plausible Pr values are the same ones that exhibit strong facilitation under control conditions (Figure 5c).

### Pronounced STP is essential for Mossy Fibers to elicit conditional detonation of CA3 Neurons

To truly grasp the functional significance of this unique STP profile, we simulated its postsynaptic consequences. To this end, we used a detailed conductance-based point model of a CA3 pyramidal neuron (CA3p) adapted from (Pinsky and Rinzel, 1994). This neuron receives synaptic input triggered by individual vesicle release events from the MF bouton. The CA3p neuron also receives many background inhibitory synaptic inputs (modeled here as constant DC input current) and exhibits low AP probability (Toth et al., 2000; Mori et al., 2004). See methods for details. However, when stimulated with a short burst of APs, mossy fibers’ fast and profound facilitation can overcome this inhibition and swiftly trigger an action potential in CA3p called detonation (Mori et al., 2004; Neubrandt et al., 2018; Henze et al., 2002). Electrophysiological studies have shown a similar response in mossy fiber synapses (Vyleta et al., 2016; Chamberland et al., 2018). This phenomenon of detonation is thought to play a crucial role in computation at this synapse by ensuring the reliable transmission of activity that is sparsely represented in granule cells—each of which makes only a few connections to CA3 neurons—while simultaneously filtering out noise. Moreover, by virtue of its binary and highly reliable postsynaptic activation, detonation enhances pattern separation within the network, increasing the orthogonality of output representations (Zalay and Bardakjian, 2006).

We observe this known sharp increase in action potential (AP) probability across successive stimuli for a subset of synaptic designs (Figure 6a). Notably, only certain control configurations reliably trigger APs in the postsynaptic CA3 pyramidal neuron. This conditional detonation is preferentially supported by synaptic designs characterized by higher values of nVDCC, dVAZ, and nAZ (Figure 6b). In stark contrast, none of the other simulation conditions consistently triggered action potentials, highlighting the unique synergy of the contributing factors of STP described in the previous section for reliable postsynaptic activation (Figure 6c, d). The probability of generating the first AP in CA3p also increases suddenly (Figure 6e). Eliminating any one of these components renders the synapse functionally ineffective. Importantly, our model uniquely reveals that collaboration between calcium domains—where calcium signals from neighboring active zones interact—is a critical and previously unknown contributor to synaptic function. Although the AP probability of CA3p increases drastically when MF is stimulated with a burst of spikes, the background inhibition dictates the delay observed in the sudden increase in AP probability (Figure 6f, g).

**Figure 6:**
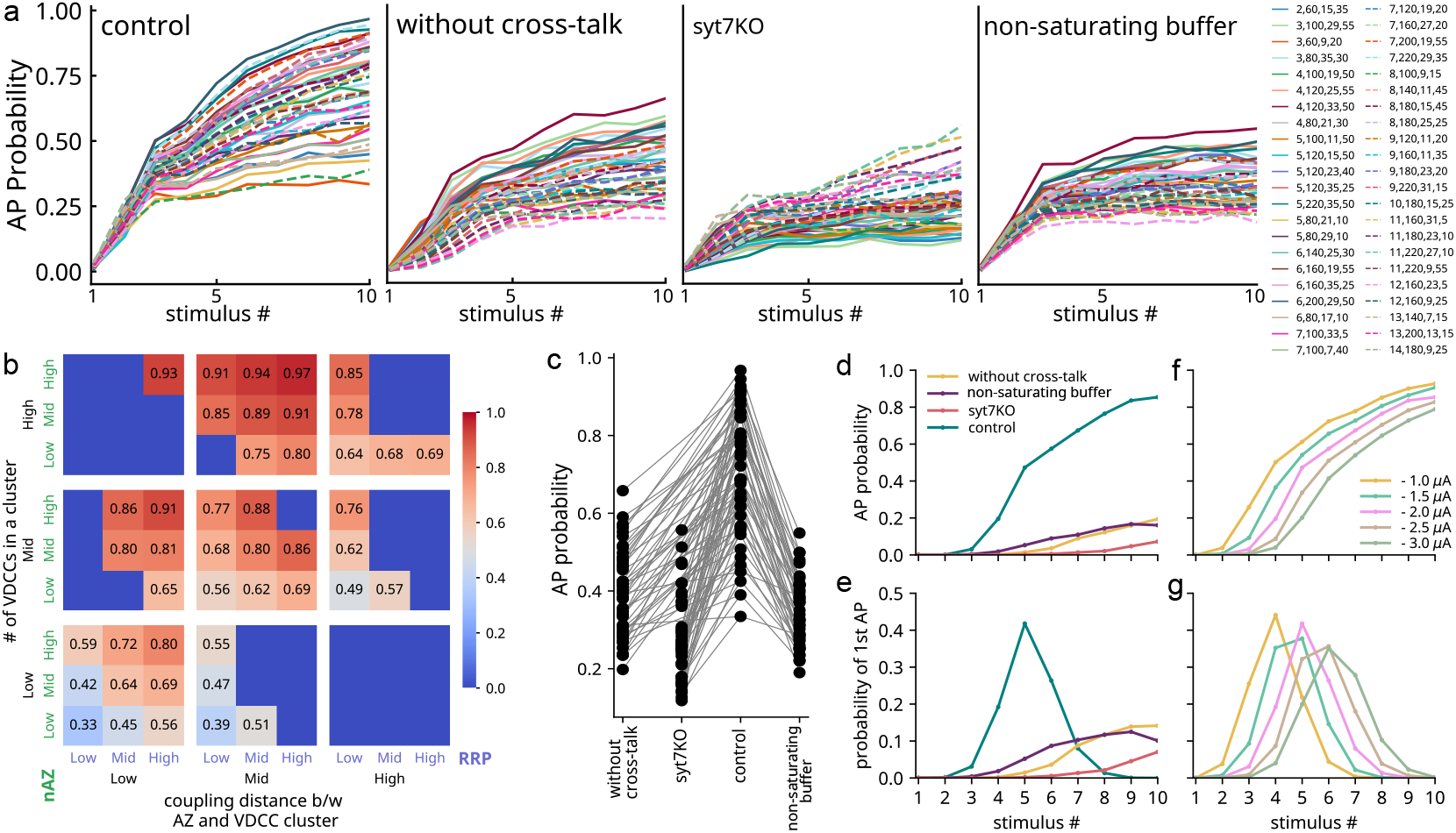
Conditional detonation of CA3 pyramidal neuron. **(a)** CA3 pyramidal neuronal model was stimulated with synaptic current generated by neurotransmitters released from mossy fiber boutons. The probability of generating an action potential in CA3p is kept low but increases drastically when stimulated with a short, high-frequency burst in control. This conditional detonation is hampered in other simulation conditions. **(b**,**c)** Different design categories show different amounts of reduction in conditional detonation, as seen here in the AP probability of CA3p after the 10th spike. **(d, e)** AP probability and probability of first AP in CA3p for different simulation conditions. **(f, g)** Delay in the sudden rise of AP probability and probability of first AP in CA3p under different inhibitory current amplitudes. The level of inhibition determines the number of presynaptic APs that CA3p requires to trigger a sudden increase in AP probability.

## Discussion

The mossy fiber to CA3 pyramidal neuron synapse is extensively studied in the mammalian brain for its role in memory and navigation. A vast body of literature details its structural complexity, molecular components, and plasticity mechanisms. Yet, despite this wealth of information, how the synapse’s intricate ultrastructure and molecular organization drive its function remains unclear. In this study, we address this fundamental question by examining the interplay between synaptic architecture, calcium buffering dynamics, residual calcium buildup, inter-AZ crosstalk, and the kinetic properties of synaptotagmin-7. Our morphologically detailed computational model offers an unprecedented opportunity to disentangle these tightly coupled factors, providing new insight into structure-function relationships that govern synaptic computation at this highly specialized synapse. Our results challenge the commonly held assumption of independent action of release sites, revealing instead the supralinear interaction between overlapping calcium domains. This synergy in calcium signaling is not an accidental feature, but a cornerstone of synaptic function at this terminal.

DG granule cells are sparsely active and fire high-frequency bursts (Neubrandt et al., 2018; Henze et al., 2002). The rapid short-term plasticity at this synapse ensures that every incoming AP contributes to a rise in vesicle release rate. CA3 neurons are depressed at basal activity levels. Only when neurotransmitter release from mossy fibers increases drastically in response to a burst does it overcome the suppression of CA3p. At this point, the probability of generating an action potential in the CA3p neuron by a single MF bouton suddenly escalates (Figure 6d). While neurons usually require multiple synapses to incrementally raise action potential probability, this synapse triggers postsynaptic spikes with exceptional potency. This allows bursts of high-frequency incoming signals to be transmitted while gating spurious spikes generated by intrinsic stochasticity. The MF bouton thus acts like a “conditional detonator,” enabling a robust transfer of input spikes arriving at the mossy fibers into information conveyed to CA3. This ability for a single MFB to trigger postsynaptic action potential also explains how sparse ‘engrams’ relay from the dentate gyrus to the CA3 region. In the subdetonation state, a single CA3 pyramidal neuron would require multiple concurrent inputs to fire. However, this scenario appears unlikely given the intrinsically sparse firing of granule cells. In contrast, under full detonation mode, activating a granule cell by a single input pattern is sufficient to elicit a postsynaptic response in CA3, providing a more efficient and plausible mechanism for sparse memory trace transmission. This phenomenon may be essential for enabling efficient encoding, storage, and retrieval of information in the granule cell–CA3 network. In summary, the exceptional short-term plasticity exhibited by this MFB underlies its specialized function. The key question that follows is: what mechanisms confer such a unique form of plasticity, distinct from any other synapse?

It turns out from our study that these large MF boutons containing many active zones with a large readily-releasable pool are optimally designed for this plasticity. A large coupling distance between active zones and their associated VDCC clusters maintains a low basal neurotransmitter release probability. Low Pr is energy efficient and provides higher tunability to the synapse. During low activity levels (spontaneous spikes), each active zone operates independently. Initially, active zones are decoupled as the spread of calcium ions is restricted due to rapid uptake by fast calcium buffers. However, during high-frequency stimuli, the cross-talk between active zones becomes significant, as the subsequent reduction in local buffering capacity allows calcium signals from one active zone to spread sufficiently to affect nearby active zones. Our study indicates that this activity-dependent initiation of cross-talk between active zones is critical for the experimentally observed rapid short-term potentiation in MF-CA3p synapses, allowing transmission of bursts.

Moreover, since synaptotagmin-7 has a low affinity for calcium (unlike synaptotagmin-1), its response to calcium influx steadily increases. Consequently, asynchronous release via synaptotagmin-7 not only exhibits a steady increase but also disperses the vesicle release over a broader time frame. This may assist in the temporal decorrelation of similar activity patterns that may represent similar experiences.

Experimental results by Vyleta et al. (2016) show that three inputs at 20 Hz can rapidly increase the probability of a postsynaptic AP; another study by (Chamberland et al., 2018) suggests six, both independent of the rate of stimuli. We found that the amount of inhibition regulates the delay in AP generation in CA3p.

Mossy fibers are under the tonic influence of adenosine, which decreases the calcium flux through VDCCs and maintains a low basal release probability in the bouton (Moore et al., 2003). Although we did not include adenosine directly into our model, its effect is accounted for by the fact that the number of VDCCs is an open parameter in our study. Therefore, lower VDCCs account for the reduced influx due to adenosine.

Complexity in the nervous system is not a product of mere numerical expansion in components or connectivity, but is tightly coordinated across multiple levels of organization, down to the molecular scale. Proteomic studies of synapses have revealed molecular complexity that correlates with both phylogenetic advancement and the increasing structural and functional complexity of the brain. Both pre- and postsynaptic terminals are increasingly understood as complex molecular systems, built from intricately organized protein networks that collectively drive emergent physiological and behavioral outcomes (Emes et al., 2008; Toth and McBain, 2000). We extend this argument beyond the synaptic proteome, recognizing that synaptic designs exhibit clear specialization across brain regions—paralleling the brain’s functional specialization. It, therefore, becomes increasingly clear that a deep understanding of synaptic designs along with their evolution is indispensable to understanding behavior itself.

## Methods

The model for the hippocampal mossy fiber bouton was built in three parts: a spatial model of the mossy fiber bouton, a stochastic point model for vesicle release via synaptotagmins (1 and 7), and a point model of the CA3 pyramidal neuron and CA3 interneuron to simulate the postsynaptic response.

### Spatial model of mossy fiber bouton

Canonical stochastic spatial model of the mossy fiber bouton has dimensions 4.1 µm × 2.5 µm × 0.8 µm (Figure 1). Active zones are placed on the synaptic surface at an inter-AZ distance of 450 nm (Rollenhagen et al., 2007). The number of active zones varies between 7 and 35. Each active zone has an associated VDCC cluster and a readily releasable pool. PMCA pumps are uniformly present on the bouton surface for slow calcium extrusion, and calbindin-D28k for buffers in the cytosol. The interplay between PMCA pumps and calcium buffers maintains a steady-state calcium concentration of 100 nm in the cytosol. MCell version 3.2.1 (https://mcell.org) was used to build and simulate the model (Stiles et al., 2001). It simulates the diffusion trajectory of each molecule within confined volumes or across surfaces, and stochastically executes user-defined reactions. Simulations were performed on a high-performance computing cluster (HP PROLIANT SL230s Gen8 as compute nodes; CPU: Intel(R) Xeon(R) CPU E5-2860 v2 *α* 2.80 GHz) housed in IISER Pune.

Voltage-dependent calcium channels are placed on the plasma membrane in small clusters 20nm across. Each cluster constitutes up to 15 channels. Depending on the simulation center-to-center distance between the active zone and its associated VDCC cluster ranges from 60 nm to 220 nm. Its kinetics are based upon Cav2.1 (P/Q-type) as characterized by Bischofberger et al. (2002). We have assumed the extracellular calcium concentration to be 2 mM for calculating calcium flux through VDCC, consistent with the experimental studies (Jonas et al., 1993; Murthy et al., 1997, 2001).

Calbindin-D28k has been modelled according to Nägerl et al. (2000). This model contains two high-affinity and two medium-affinity calcium binding sites. The bouton is populated with calbindin-D28k at a concentration of 40 µM (Muller et al., 2005). The simulations with unsaturating buffers are implemented by assuming a constant buffer concentration and multiplying it by the buffer rate constants.

PMCA is present on the plasma membrane with a density of 180 µm^−2^. The kinetic model for the PMCA pump has been taken from Penheiter et al. (2003); Brini and Carafoli (2009) and implicitly incorporates calcium leak across the plasma membrane.

The model for calcium sensors (synaptotagmin 1 and 7) for neurotransmitter release at MFB has been taken from our previous work (Nadkarni et al., 2010). It is implemented using STochastic Engine for Pathway Simulation (STEPS) version 3.5.0 (https://github.com/CNS-OIST/STEPS), a Python package for exact stochastic simulation of reaction-diffusion systems using Gillespie’s SSA (Hepburn et al., 2012). This model incorporates calcium-dependent fast synchronous and slow asynchronous, and calcium-independent spontaneous modes of vesicle release. Local calcium concentrations at each active zone—measured from the MCell model—are fed to the system, and individual neurotransmitter releases are recorded.

Kinetic schemes used for each of the components and the corresponding reaction rates are taken from our previous study (Singh et al., 2021).

### CA3 pyramidal neuron model

CA3 neurons were based on the model by Pinsky and Rinzel (1994). In addition to the Pinsky-Rinzel model, synaptic currents (AMPA and NMDA) were implemented in response to neurotransmitter release from mossy fiber bouton (Equation 2, Equation 3, parameters: table Table 2). Input from the presynaptic terminal is fed to the CA3 pyramidal neuron model as synaptic currents. The release of neurotransmitter updates AMPA and NMDA currents via Equation 4. We assumed each active zone has an associated independent cluster of AMPA receptors on the postsynaptic terminal. Brian version 2.9.0 (https://briansimulator.org), a Python package, was used to implement the model (Stimberg et al., 2019).

**Table 2.** Model Parameters.

| Parameter | Value |
| --- | --- |
| $\tau_{\text{AMPA}}$ | 5.0 ms |
| $g_{\text{AMPA}}$ | 0.2 mS |
| $V_{\text{AMPA}}$ | 0.0 mV |
| $g_{\text{NMDA}}$ | 0.1 mS |
| $V_{\text{NMDA}}$ | 0.0 mV |
| $\tau_{\text{NMDArise}}$ | 2.0 ms |
| $\tau_{\text{NMDAdecay}}$ | 100.0 ms |
| $\alpha$ | 0.5 ms<sup>-1</sup> |
| $\text{Mg}^{2+}$ | 1.0 |

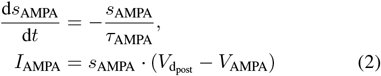

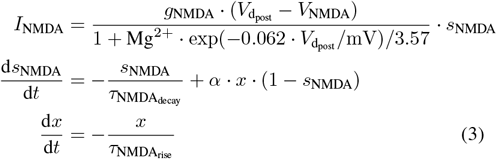

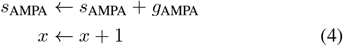

## Acknowledgements

This work was supported by Pratiksha Trust (EMSTAR/22/078) and the Indian Institute of Science Education and Research, Pune.

## Author Contributions

N.S.: conceptualization, methodology, software, validation, formal analysis, investigation, data curation, writing original draft, and visualization. S.N.: conceptualization, methodology, software, validation, formal analysis, resources, investigation, writing original draft, reviewing, editing, visualization, supervision, project administration, and funding acquisition.

## Code Availability

The code used in this study is available at https://github.com/nishs1729/MFB

